# A standardized broad host range inverter package for genetic circuitry design in Gram-negative bacteria

**DOI:** 10.1101/2020.07.14.202754

**Authors:** Huseyin Tas, Ángel Goñi-Moreno, Víctor de Lorenzo

**Affiliations:** Systems Biology Department, Centro Nacional de Biotecnología-CSIC, Campus de Cantoblanco, Madrid 28049, Spain; School of Computing, Newcastle University, Newcastle Upon Tyne, NE4 5TG, UK; Centro de Biotecnología y Genómica de Plantas (CBGP, UPM-INIA), Universidad Politénica de Madrid (UPM), Instituto Nacional de Investigación y Tecnología Agraria y Alimentaria (INIA), Campus de Montegancedo-UPM, 28223 Pozuelo de Alarcón, Madrid, Spain

**Keywords:** Broad host range tools, inverter library, gates, Pseudomonas putida, flow cytometry, biosensor, automated circuit design

## Abstract

Genetically encoded logic gates, especially inverters—NOT gates—are the building blocks for designing circuits, engineering biosensors or decision-making devices in synthetic biology. However, the repertoire of inverters readily available for different species is rather limited. In this work, a large whole of NOT gates that was shown to function previously in a specific strain of *Escherichia coli*, was recreated as broad host range (BHR) collection of constructs assembled in low, medium and high copy number plasmid backbones of the SEVA (Standard European Vector Architecture) collection. The input/output function of each of the gates was characterized and parameterized in the environmental bacterium and metabolic engineering chassis *Pseudomonas putida*. Comparisons of the resulting fluorescence cytometry data with those published for the same gates in *Escherichia coli* provided useful hints on the portability of the corresponding gates. The hereby described BHR inverter package (20 different versions of 12 distinct gates) thus becomes a toolbox of choice for designing genetic circuitries in a variety of Gram-negative species other than *E. coli.*

---

Design and implementation of genetic circuits is one of trademarks of contemporary synthetic biology^*1*^. The archetypal approach involves abstracting biological cues (e.g. effectors, metabolites, proteins) and physicochemical signals (e.g. inducers, temperature) as inputs. These are processed through a more or less complex computation layer (most often assembled with regulatory parts: transcriptional factors, riboregulators etc.) which then releases another biological signal as the output. This in turn can be used as the input for another node of the circuit and so on^*2, 3*^. Further abstraction of biological circuits as wholes of Boolean logic gates enables a superior level of complexity, as shown by a suite of examples involving rewiring of stress responses, detection of environmental contaminants, implementation of cellular calculators and others^*4–7*^. Yet, as the demand for increasingly complex circuits grows^*5, 8, 9*^, so does the interest in automation of their design and execution. One major landmark in this direction was the publication in 2016 of the CelloCAD platform^*5*^, a complete operative system for virtual assembly and eventual implementation of logic circuits in *E. coli* through combination of series of NOR gates—based themselves on a large collection of well characterized promoter/repressor pairs (i.e. inverters or NOT gates). This tool affords automated design and simulation in seconds of complex genetic networks with successful prediction of > 70% of all states shown by the actual DNA constructs once synthesized and knocked in *E. coli^5, 7^*. Yet, due to the material nature of the building blocks, CAD tools such as Cello are inherently restrictive to single strains or species. While these systems may work well in a given organism, the parameters and general behavior can change very significantly when passed to other hosts. This is a considerable issue when circuits are desired to compute signals under environmental and industrial conditions for which *E. coli* is not an optimal chassis.

On this basis we set out to engineer a robust and easy-to-use package of standardized NOT gates that could be used for circuit design in Gram-negative bacteria other than *E. coli* and that—once characterized in the host of interest—could benefit from the CelloCAD software^*5*^ for automatic assembly of the cognate DNA.

To this end, we recreated the DNA sequence encoding each of the inverters available in the Cello platform and pass them to vectors of the broad host range SEVA (Standard European Vector Architecture) collection with low, medium and high copy numbers. The general organization of the constructs is sketched in Fig. 1. Note that the business part of each plasmid is flanked by upstream and downstream terminators added by the SEVA structure to mitigate potential readthrough from vector promoters. The plasmids backbones were retrieved from the SEVA database and repository^*10*^.

**Figure 1.**
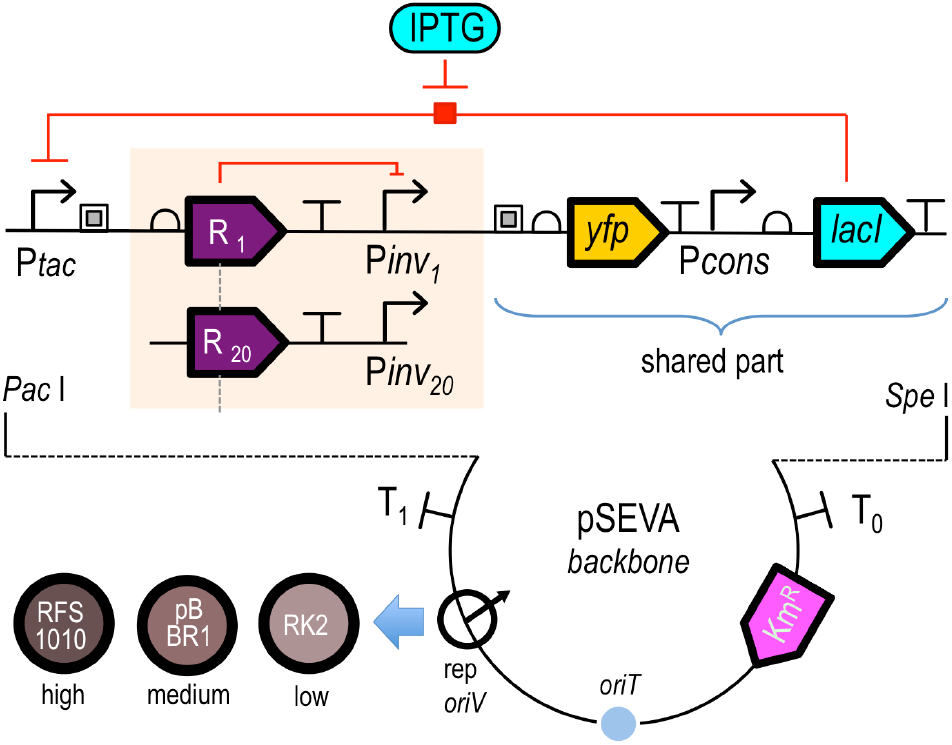
Schematic representation of BHR genetically-encoded inverters. The organization of the functional parts borne by the 20 variants assembled following the SEVA standard is sketched (components not to scale). Similarly to the earlier collection for *E. coli^5^* there are segments shared by all construct i.e. the P*tac/lacI*/IPTG-dependent expression system for each of the repressor/SD combinations (R_1_ to R_20_) and the *yfp* gene used for fluorescent readout of inverter performance (the *lacI* gene is expressed through the same constitutive promoter P*cons*). The Km resistance gene and the DNA sequences that punctuate the SEVA backbone (T_1_ and T_0_ terminators, origin of conjugal transfer *oriT*) are common to all constructs as well. The variable parts include [i] the DNA encoding the SD, the repressor gene (R) and the cognate repressible promoter (P*inv*) upstream of the reporter *yfp* and [ii] the BHR origin of replication: RK2, pBBR1 or RSF1010, each of them supporting different plasmid copy numbers as indicated.

In practical terms, each of the segments encoding the gates was amplified from the collection of *E. coli* NEB10β strains bearing p15A/Km^R^ vectors (~15 copies) inserted with the cognate DNA^*5*^. Primers (Merck Sigma Aldrich, Inc) were designed for adding SEVA-compatible PacI and SpeI restriction sites to the extremes of the amplicons generated with the Q5 High-Fidelity DNA polymerase (New England BioLabs, Inc.). The oligonucleotides used for amplification of the DNA of the NOT gates and verification of the constructs are listed in Supplementary Table S2. Following PCR, the resulting fragments where separately cloned in pSEVA221 (RK2*oriV*, ≤ 5 copies/cell), pSEVA231 (BBR1*oriV*, ~ 30 copies) and pSEVA251 (RFS1010*oriV*, ≥ 50 copies) and first captured, verified and resequenced in the host strain *P. putida* KT2440. The complete catalogue of constructs is listed in Table 1.

**Table 1.** Library of available genetically-encoded inverters and auxiliary plasmids in SEVA vectors with 3 different copy numbers (see Supplementary Table S1 for plasmid names and cargoes). Note that 5 inverters caused growth inhibition when cloned in high copy number vectors, plausibly due to toxicity of the repressor protein encoded therein. The repressor borne by each of the gates is described in^*5*^.

| Inverters | Low Copy Number | Medium Copy Number | High Copy Number |
| --- | --- | --- | --- |
| AmeR-F1 | ✓ | ✓ | ✓ |
| AmtR-A1 | ✓ | ✓ | ✓ |
| BetI-E1 | ✓ | ✓ | ✓ |
| BM3R1-B1 | ✓ | ✓ | - |
| BM3R1-B2 | ✓ | ✓ | ✓ |
| BM3R1-B3 | ✓ | ✓ | ✓ |
| HlyIIR-H1 | ✓ | ✓ | ✓ |
| IcaRA-I1 | ✓ | ✓ | ✓ |
| LitR-L1 | ✓ | ✓ | ✓ |
| LmrA-N1 | ✓ | ✓ | ✓ |
| PhlF-P1 | ✓ | ✓ | ✓ |
| PhlF-P2 | ✓ | ✓ | ✓ |
| PhlF-P3 | ✓ | ✓ | - |
| PsrA-R1 | ✓ | ✓ | ✓ |
| QacR-Q1 | ✓ | ✓ | ✓ |
| QacR-Q2 | ✓ | ✓ | - |
| SrpR-S1 | ✓ | ✓ | ✓ |
| SrpR-S2 | ✓ | ✓ | - |
| SrpR-S3 | ✓ | ✓ | - |
| SrpR-S4 | ✓ | ✓ | ✓ |
| Auto-fluorescence | ✓ | ✓ | ✓ |
| Standardization | ✓ | ✓ | ✓ |
| Promoter Activity | ✓ | ✓ | ✓ |

Note that some gates (those designated BM3R1-B1, PhlF-P3, QacR-Q2, SrpR-S2 and SrpR-S3) could not be cloned in the higher copy number vectors, presumably

Note that some gates (those designated BM3R1-B1, PhIF-P3, QacR-Q2, SrpR-S2 and SrpR-S3) could not be cloned in the higher copy number vectors, presumably due to the toxicity effects of the encoded repressor proteins. The collection (Table 1 and Supplementary Table S1) includes a total of 12 inverters, some of them bearing two or more variants of the Shine-Dalgarno sequence of the repressor gene that changes its expression levels. For example, AmeR-F1 (AmeR is the gate and F1 is the SD) has only 1 version of the ribosomal binding site of the repressor gene; however, SrpR has 4 versions (SrpR-S1, SrpR-S2, SrpR-S3, SrpR-S4 (SrpR is the gate and S1, S2, S3 and S4 the 4 different SDs)). Note also that the arrangement of functional parts of the gates in the SEVA vectors is identical to the one originally adopted in the p15A/Km^R^ vectors of the Cello platform (Fig. 1). In addition to the plasmids bearing 20 inverters, we built a set of reference constructs allowing relative promoter units (RPU) to be converted and compared between different conditions (see below). These references (Supplementary Fig. S1) include [i] autofluorescence plasmids (recreating the business parts of pAN1201^*5*^ and consisting of each of the backbone plasmids but without any insert e.g. missing the repressor/target promoter segments highlighted in Fig. 1, [ii] RPU standard plasmids derived from pAN1717^*5*^ in which *yfp* is expressed through the constitutive promoter J23101 and which are used in combination with the autofluorescence plasmid for converting YFP readouts of each inverter into RPU (see equation #1 below) and [iii] promoter activity plasmid recreating pAN1818^*5*^, which is used for measuring the promoter activity (pTac-YFP) based on inducer concentrations (Supplementary Fig. S1).

The library of constructs listed in Table 1 and Supplementary Table S1 is expected to ease utilization of CelloCAD as a genetic programming tool in a suite of Gram-negative bacteria. Note that users have a choice to pick the same gate borne by plasmids with different copy numbers, what may be critical to avoid potential toxicity. Adoption of the standard SEVA format gives two additional advantages. While the construct library of Supplementary Table S1 was built on vectors with a Km resistance gene, the modularity of the SEVA format makes its replacement by an alternative antibiotic marker for selection^*10, 11*^ easy. Also, the BHR nature of the standardized constructs affords their implantation in diverse Gram-negative hosts, thereby giving a chance to compare their performance in different species and thus learn about interoperability and context dependencies in circuit designs.

In order to validate these features we tested and parameterized the whole low-copy number versions of the inverter library of Supplementary Table S1 in the environmental bacterium and Synthetic Biology chassis *Pseudomonas putida* KT2440^*12, 13*^. The experimental workflow to this end (Fig. 2) was designed considering the specific needs of *P. putida* for growth as detailed in the legend of the figure. Once strains bearing each of the constructs were generated, transformants were grown and IPTG concentrations of (5-1000 μM) added to activate each of the devices, for a total 24 h period. YFP fluorescence emission detected with a flow cytometer was recorded after 24 of IPTG addition. Data were then analyzed with FlowJo software (https://www.flowjo.com/). An important detail was that the auto-gating option of the software was set considering at least 50% of the events covered while Forward and Side scatters were plotted. The same gating conditions were applied to all specimens in the same group and repeated for all the samples.

**Figure 2.**
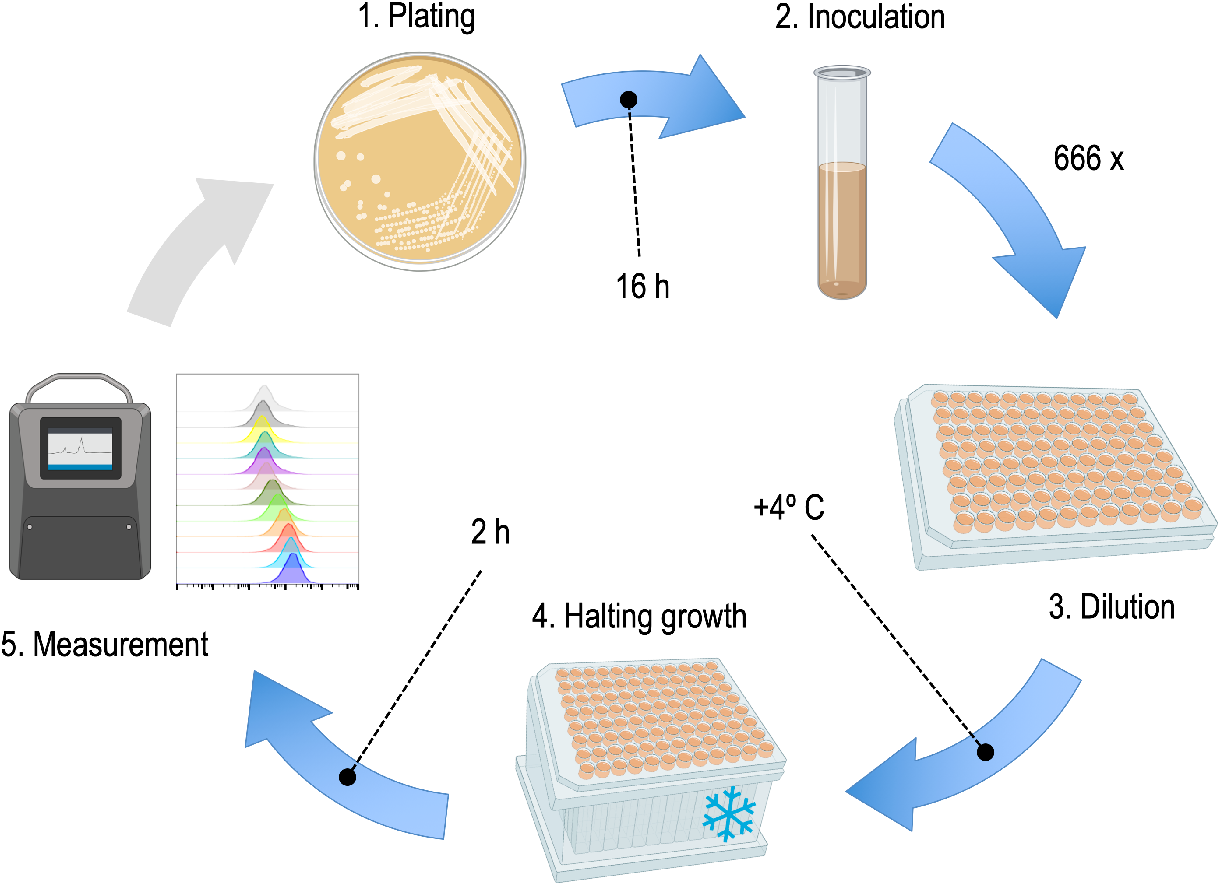
Experimental protocol used for measuring performance of the genetic inverters listed in Table 1 in P. putida KT2440. In all experiments bacteria were grown in M9 medium adapted for growth of *P. putida*. 250 ml of liquid culture contained 25 ml x10 M9 salts, 500 μl of 1M MgSO_4_, 2.2 ml of 20% citrate and milliQ-H_2_O to volume. 50 μg ml^−1^ kanamycin was also added to secure plasmid retention (all them Km^R^). For the experiments, a saturated overnight culture of each of the strains under were diluted over ~ 600-fold in the wells of a microtiter plate with 200 μl per well, plated for 24h, added with IPTG concentrations ranging 0 to 1000 μM and incubated at 30 °C with shaking for 24 h. Cultures (typically reaching OD_600_ ~ 0.2-0.3) were then kept in the cold for the rest of the procedure. YFP fluorescence distribution of each sample was measured with a Miltenyi Biotec MACS flow cytometer at channel B1 with an excitation of 488 nm and emission of 525/50 nm. For each sample 30 thousand events were collected with singlet gating. Calibration was done by using MACSQuant Calibration Beads (see text for explanation).

On the basis of the thereby produced data we quantified the output of the individual devices at each condition as standard RPUs (relative promoter units). This was done by characterizing the fluorescence values emitted by the bacterial population of the cultures exposed to IPTG levels covering an induction ranges from none to saturation (in our case 12 points of growing effector concentrations). On this basis, the RPU value at each point can be calculated with the formula (1):

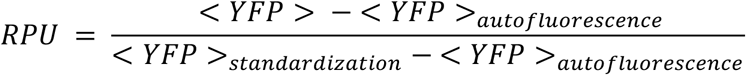

Where <YPF> is the median fluorescence value from the gate of interest that is to be converted into RPU (from either the inverter-bearing plasmids or the control promoter activity plasmid), <YPF>_autofluorescence_ is the median fluorescence value of auto-fluorescence plasmid, <YPF>_standardization_ is the median fluorescence value from the standardization plasmid. Next, the output *vs* input values of each inverter were represented in a response function plot generated with Hill fits and utilizing the RPU figures as the input to the calculations. Specifically, Hill equation parameters were estimated by plotting each inducer levels with their corresponding standardized median florescence values (Supplementary Data S2) by means of the equation (2):

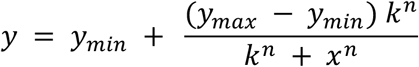

where *y* is the output promoter activity, *y_min_* is the minimum observed value for promoter activity, *y_max_* is the maximum observed value for promoter activity, *k* is the input value for which half maximum value for output promoter is reached, *n* is the Hill coefficient. Experimental response data were thus entered in the above Hill equation. The corresponding Hill parameters are provided in Supplementary Table S3 for both *E. coli* (retrieved from www.cellocad.org) and *P. putida* values (this study). Fitting was performed with MATLAB using the scripts provided in Supplementary Data S1.

The resulting characterization of the 20 inverters (listed in the Supplementary Table S1) in *P. putida* based on the protocols and calculations above is shown in Fig. 3. The data showed a range of divergences in the behavior of the inverters in either host. In some cases, the patterns were comparable both in terms of the dynamic ranges of the input/output, the contour of the response curves and the specific values of promoter strengths. In other cases, the shape of the curve was kept but the boundaries changed very significantly. And finally, in yet another series of devices there was little if any similarity in their input/output transfer functions between the two types of bacteria While these data expose the limitations of just exporting genetic devices from one species to the other, the large repertoire of gates also enable users to pick the ones whose parameters are compatible with the CelloCAD tool for automation of circuit designs^*5*^. Furthermore, the dataset associated to Fig. 3 encrypts valuable, quantitative information on the interoperability of parts and circuits between different biological recipients of the same constructs, an issue hardly tackled thus far in the Synthetic Biology literature^*14, 15, 16*^

**Figure 3.**
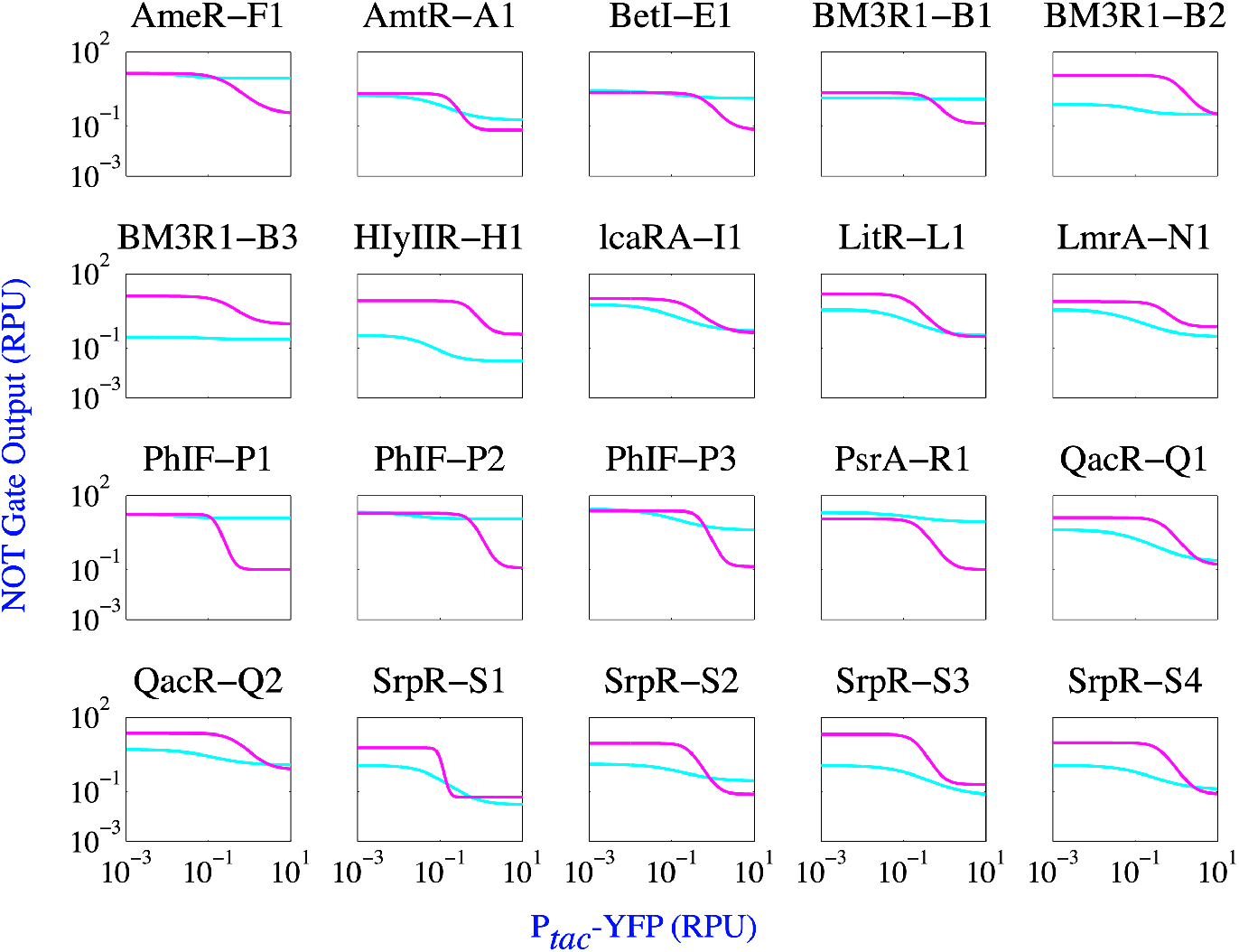
Behavior of genetically-encoded inverters in *P. putida* vs. *E. coli.* The panel shows the comparisons of the of the Hill of the same inverters in *E. coli* NEB10β (magenta; data retrieved from^*5*^) and behavior the same devices in *P. putida* KT2440 (cyan; experimental from this study). X-axes correspond to the activity of the IPTG-inducible pTac promoter and Y-axes indicate the activity of corresponding inverters. Both axes indicate YFP expression in RPUs.

In sum, we believe that the hereby described collection of inverters available in a BHR format will help to expand the possibilities of genetic programming towards bacteria other than *E. coli* but still interesting from a SynBio-inspired biotechnological perspective. Furthermore, their testing and parameterization in various hosts may deliver general portability rules that thus far rely on a mere trial-and-error exercise. The whole collection of constructs is available through the SEVA database and vector repository at http://seva.cnb.csic.es.

## Associated content

## Supporting information

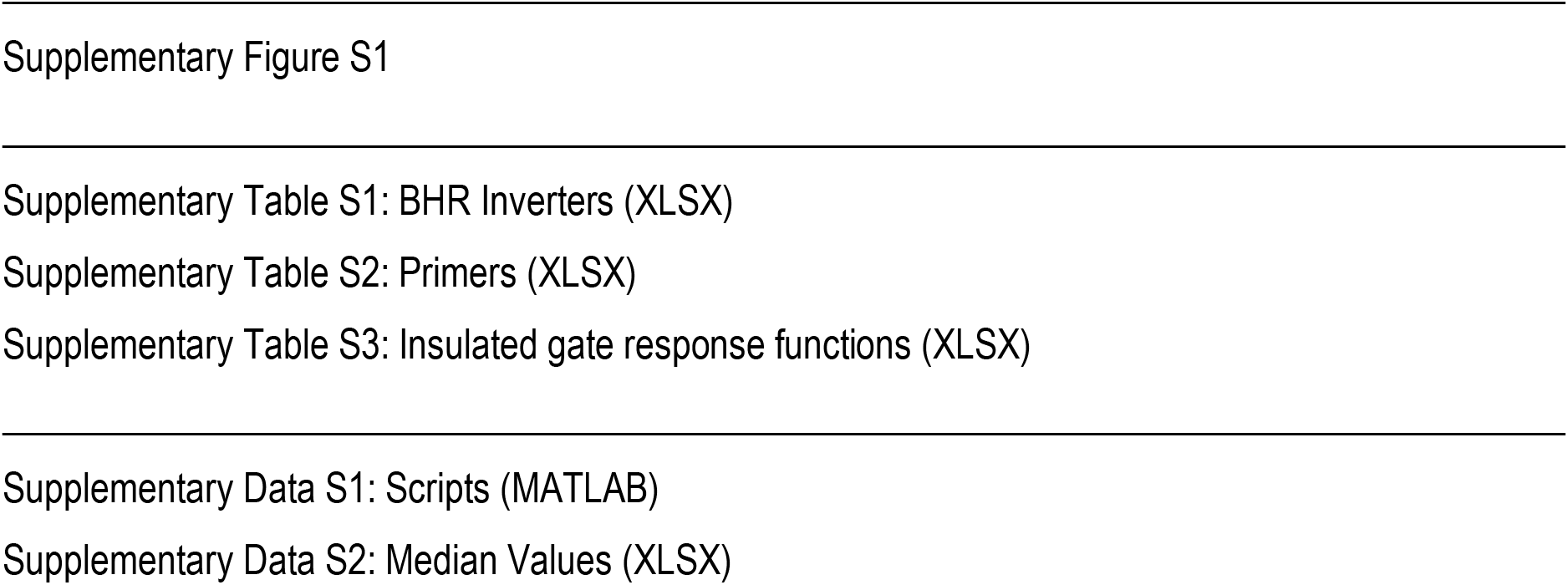

## Acknowledgements

We are indebted to Chris Voigt and Alec Nielsen (MIT) fort sharing the collection of NOT gates developed for the Cello platform.

## Author Contributions

HT, AG-M and VdL planned the experiments, analyzed and discussed the data and contributed to the writing of the article.

## Funding

This work was funded by the SETH Project of the Spanish Ministry of Science RTI 2018-095584-B-C42, the MADONNA (H2020-FET-OPEN-RIA-2017-1-766975), BioRoboost (H2020-NMBP-BIO-CSA-2018), SYNBIO4FLAV (H2020-NMBP/0500) and MIX-UP (H2020-Grant 870294) Contracts of the European Union, the S2017/BMD-3691 InGEMICS-CM Project of the Comunidad de Madrid (European Structural and Investment Funds) and the SynBio3D project of the UK Engineering and Physical Sciences Research Council (EP/R019002/1).

## Notes

The authors declare no competing financial interest

## Abbreviations

BHR: broad host range
SD: Shine-Dalgarno sequence
RBS: ribosomal binding sequence
IPTG: isopropyl thiogalactopyranoside
CAD: computer-assisted design
SEVA: Standard European Vector Architecture

## Notes

### Competing Interest Statement

The authors have declared no competing interest.

