## Supplementary Fig S1 for "A standardized broad host range inverter package for genetic circuitry design in Gram-negative bacteria"

TECHNICAL NOTE

by Huseyin Tas, Angel Goñi-Moreno and Víctor de Lorenzo

Supplementary **Figure S1.** Plasmids of reference for calculation of relative promoter units (RPU)

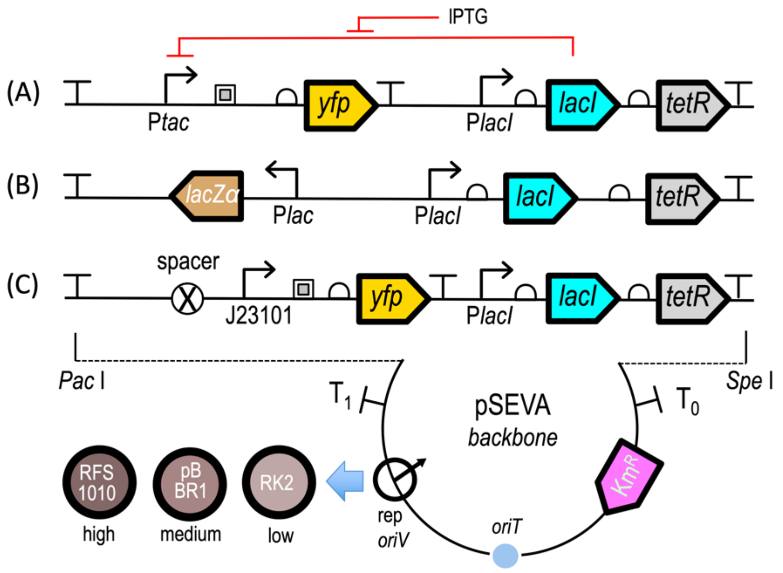


(A) Promoter activity plasmid: sensors LacI and TetR are encoded in the same operon at the downstream of PlacI constitutive promoter. The sensors are to be used with corresponding promoter in this case Ptac is used. Ptac is regulated by LacI expressed in the operon and by the inducer IPTG. In between Ptac and *yfp* gene RiboJ ribozyme is used as a promoter context insulator (square glyph). By regulating IPTG levels the activity of the Ptac promoter is measured by YFP levels. The SEVA backbone terminators ($T_0$ and $T_1$) are kept to insulate the circuit encoded in-between. $Km^R$ is the selection markers in all cases. The BHR origin of replication was prepared in 3 versions: RK2, pBBR1, RFS1010.

(B) Autofluorescence plasmid: This construct is used as the backbone of all constructs as published previously [1]. Sensors LacI and TetR are encoded in the same operon at the downstream of PlacI constitutive promoter. lacZa gene is expressed by constitutive Plac promoter. The terminator at the downstream of Plac is L3S2P21 terminator and tetR gene is insulated with native AraC terminator.

(C) RPU standard plasmid. Sensors LacI and TetR are encoded in the same operon at the downstream of PlacI constitutive promoter. yfp gene is expressed by standard constitutive J23101 promoter. There is a 15 bp spacer upstream of the J23101 promoter, and a terminator upstream of it. Another terminator (L3S2P21) is located downstream of yfp. Finally, tetR gene is insulated with native AraC terminator.

[1] Nielsen, A. A., Der, B. S., Shin, J., Vaidyanathan, P., Paralanov, V., Strychalski, E. A., Ross, D., Densmore, D., and Voigt, C. A. (2016) Genetic circuit design automation, Science 352, aac7341.
