## Supplementary Data S1 for "A standardized broad host range inverter package for genetic circuitry design in Gram-negative bacteria": fitting.m

```
function entries = fitting(Data, labels)

gateX = Data(:,1);

entries = [];
for i = 1:20

    gateY = Data(:,i+1);
    [params, score] = fitting_gate(gateX, gateY);
    % params is the parameters
    % score is the good of fit
    entry = [labels(i+1), score, params(1), params(2)];
    entries =[entries; entry];
   
end
```
