## Supplementary Data S1 for "A standardized broad host range inverter package for genetic circuitry design in Gram-negative bacteria": fitting_gate.m

```
function [minimizer, score] = fitting_gate(x_array, gate_array)
    
    objective_function = @(p) loss_fun(p,x_array, gate_array);
    
    initial_p = [0.09 1.4];
    lb = [0, 1];
    ub = [Inf, Inf];
    
    [minimizer, score] = fmincon(objective_function, initial_p, [], [], [], [], lb, ub);
        
end
```
