## Supplementary Data S1 for "A standardized broad host range inverter package for genetic circuitry design in Gram-negative bacteria": mean_logsqerr_fun.m

```
function e = mean_logsqerr_fun( actual, predicted )

%This function does "MEAN SQUARED LOGARITHMIC ERROR" _ Array Version

    abs_e = arrayfun(@(a,p) logsqerr(a, p), actual, predicted);

    e = mean(abs_e);


end
```
