## Supplementary Data S1 for "A standardized broad host range inverter package for genetic circuitry design in Gram-negative bacteria": plotting.m

```
function entries = plotting(Data, Data2, labels)

Data = load('Data.mat');
Data2 = load('Data2.mat');
labels = load('labels.mat');
Data = cell2mat(struct2cell(Data));
Data2 = cell2mat(struct2cell(Data2));
labels = struct2array(labels);
entries = fitting(Data, labels);
entries2 = fitting(Data2, labels);

gateX = Data (:,1);
gateX2 = Data2 (:,1);
figure
for i = 2:21
    gateY = Data(:,i);
    gateY2 = Data2(:,i);
    subplot(4,5,i-1);

  
    parameter = entries(i-1,3:4);
    parameter = cell2mat(parameter);
    hill_curve = hill(min(gateY),max(gateY),parameter(1),parameter(2));
    
   
    xs = 10.^[-3:0.01:1];
    ys = arrayfun(@(x) hill_curve(x), xs);
    loglog(xs,ys,'Linewidth',2,'Color','c');
    hold on

    
    parameter2 = entries2(i-1,3:4);
    parameter2 = cell2mat(parameter2);
    hill_curve2 = hill(min(gateY2),max(gateY2),parameter2(1),parameter2(2));
    
    xs2 = 10.^[-3:0.01:1];
    ys2 = arrayfun(@(x2) hill_curve2(x2), xs2);
    loglog(xs2,ys2,'Linewidth',2,'Color','m'); 
    title(labels(i), 'FontSize', 24, 'FontWeight', 'bold');
    hold on
   
    
    set(gca, 'XLim',[0.001 10]);
    set(gca, 'YLim',[0.001 100]);
    
    set(gca,'xscale','log')
    set(gca,'yscale','log')
    
    set(gca,'xtick', [0.001 0.1 10]);
    set(gca,'ytick', [0.001 0.1 100]);
    set(gca,'fontsize',16, 'fontname', 'times', 'fontweight', 'bold');

    
    hold off
    
end
end
```
